# Assembly Mechanisms of Neotropical Butterfly Communities Along an Environmental Gradient

**DOI:** 10.1101/632067

**Authors:** María F. Checa, David Donoso, Elisa Levy, Sebastián Mena, Jaqueline Rodriguez, Keith Willmott

**Author notes:** **Correspondence:** Maria F. Checa, Museo QCAZ de Invertebrados, Pontificia Universidad Católica del Ecuador, Apartado Postal 17-01-21-84, Quito, Ecuador.

## Abstract

Our goal was to test the hypothesis that assembly processes that limit species similarity (i.e., competition) predominantly occur in more ‘stable’ abiotic environments, whereas habitat filtering (i.e., habitat characteristics) is a major driver of community composition within more variable environments at regional (e.g., aseasonal vs seasonal forests) and local scales (e.g., understory vs. canopy). A combined approach of phylogenetic- and functional trait-based analyses using forewing length and aspect ratio as traits, were used to this hypothesis.

A 3-year survey was carried out at three sites (i.e., wet, transition and dry forests) across a climatic gradient in western Ecuador. Transition and dry forests were considered as seasonal, whereas wet forest were considered aseasonal. Butterflies were sampled using traps baited with rotting banana and prawn every two months from Nov 2010 to Sep 2013. Traps were set up at two heights, in the understory and canopy. DNA was extracted to sequence the barcode’ section of the mitochondrial gene cytochrome oxidase 1 (COI) for phylogenetic analyses. Measurements of morphological traits, forewing length and aspect ratio were done using digital photographs of specimens.

A total of 6466 specimens representing 142 species of Nymphalidae were recorded. Based on phylogenetic- and trait-based analyses, we rejected the hypothesis that assembly processes that limit species similarity (i.e., competition) are likely to predominantly occur in more ‘stable’ abiotic environments, whereas habitat filtering can be a major driver of community composition within more variable environments at regional (i.e., aseasonal forest vs seasonal forests) and local scales (i.e., understory vs. canopy). My study of assembly mechanisms revealed the opposite pattern, with stronger evidence for the action of ecological filters in the assembly of butterfly communities from the wet aseasonal forests, and competition likely to be a major assembly process within dry seasonal forests. The present study therefore provided new insights into community assembly mechanisms in one of the richest butterfly faunas worldwide.

## Introduction

Spatial variation in species richness is one of the most obvious attributes of biological communities, but the causes of such variation remain one of the most actively studied and debated areas in ecology (Mittelbach 2012). Species richness peaks in equatorial regions and gradually decreases towards the poles for virtually all taxonomic groups; several hypotheses have been proposed to explain this gradient including ecological, biogeographic and evolutionary processes as determinant factors (Schemske *et al.* 2009, Wiens & Donoghue 2004, Valentine *et al.* 2008, Schemske 2009).

A majority of studies have shown a correlation between environmental factors and diversity patterns, with the energy hypothesis being one of the most broadly accepted hypothesis to explain these patterns (see Tello & Stevens 2010). Climate is hypothesized to play a key role driving the latitudinal pattern of species richness through energy availability, measured as primary productivity (Williams & Middleton 2008, Williams & Hero 2001, Hanya *et al.* 2011). Evidence supporting the energy hypothesis has been provided by studies with birds (Evans *et al.* 2006, Williams & Middleton 2008), bats (Tello & Stevens 2010) and primates (Hanya *et al.* 2011). Climatic variables are frequently used as reasonable surrogates of primary productivity, but seasonality of climate can likely better predict species richness than total or mean measurements (Pyrcz *et al.* 2014), particularly in seasonal forests (Williams & Middleton 2008). Areas with high seasonal rain variation have lower species richness of frogs (Williams & Hero 2001) and birds (Williams & Middleton 2008). Hanya *et al.* (2011) found that species richness increased as seasonality of resource availability decreased and the effects of biogeographic factors, if any, were small, revealing the importance of seasonality in determining energy availability.

Despite strong correlations between climate and species diversity, however, the mechanisms by which climate controls diversity are more poorly understood. Resource-niche partitioning is hypothesized to promote greater species diversity (Finke & Snyder 2008) since it decreases interspecific competition, a condition denoted in early ecological models (Hutchinson & MacArthur 1959, MacArthur & Levins 1967, Chase & Leibold 2003) to foster species coexistence and promote biodiversity. One possibility is therefore that more stable climates, which host higher species diversity permit greater niche specialization as a result of continual competition, allowing more species to coexist, whereas less stable climates favor broader niches, that perhaps require specific adaptations, and thus result in less diverse communities. Phylogenetic and phenotypic analyses are an increasingly common approach to examine the relative importance of these two kinds of community assembly mechanisms, namely competition and habitat filtering (Mittelbach 2012). These analytical tools allow inferring about the action of niche-based processes in community assembly by examining patterns of phylogenetic and trait dispersion within communities (Cavender-Bares *et al.* 2009, Donoso 2013). The distribution of traits and phylogenetic distance observed within a community is then compared to ‘expected’ values generated within null communities formed by randomly selecting species from a regional pool of potential colonists (Webb 2000, Kraft *et al.* 2007).

One of the most familiar stabilizing mechanisms (*i.e.*, processes that give advantage to species when rare) that occur in tropical forests is niche partitioning with respect to heterogenous light environments across different habitats (Harms 2001) and strata (Terborgh 1985), which is expected to generate communities with co-occurring species sharing adaptations to the environment, consisting in traits to use shared resources (Kraft *et al.* 2010). This mechanism along with other examples encompassing different habitat characteristics are referred to as habitat filtering, a process that influences the range of ‘viable’ trait values existing at a given site (see Cornwell & Ackerly 2009).

On other hand, competition influences the spacing of trait values within a community, imposing limits to the similarity of coexisting species, a prediction of classical models of competition (MacArthur & Levins 1967, Cornwell & Ackerly 2009). For traits exhibiting phylogenetical signal (*i.e.*, phylogenetic conservatism), ecological filters result in communities formed of closely related species (*i.e.*, phylogenetic clustering) (Kraft *et al.* 2007, Lessard *et al.* 2009, Kraft *et al.* 2010), whereas competition produces an even dispersion of traits and co-occurring species are less related than expected (Webb 2000). With respect to convergent traits, assembly processes can generate completely opposite patterns of community phylogenetic dispersion (Kraft *et al.* 2007). Furthermore, if traits are irrelevant as drivers of community assembly, a precept of neutral theory (Hubbell 2001), random patterns of trait and phylogenetic dispersion are expected outcomes.

The relative importance of habitat filtering and competition might differ along environmental gradients. A study of hummingbird communities in Ecuador revealed an interesting pattern: competition constituted the dominant mechanism driving community assembly within moist lowland forests, whereas communities inhabiting seasonally dry forests were predominantly shaped by habitat filtering (Graham *et al.* 2009). Likewise, similar phylogenetic clustering within ‘harsh’ environments compared to more ‘benign’ conditions has been observed for plant traits across latitude and/or elevation (Sweson & Enquist 2007, Cornwell & Ackerly 2009). Nevertheless, it is unknown whether these patterns apply to other biological communities such as tropical insects, as studies focused on assembly mechanisms of tropical communities remain scarce. This is particularly true for insects (Chazot *et al.* 2014, Donoso 2013) along environmental gradients, such as wet moist lowland to dry forests. Nevertheless, studies of insects have great potential, since insects are the richest group of living organisms (Wiens *et al.* 2015), concentrated in the tropics, with nearly half of the world’s species inhabiting these forests (Olson & Dinerstein 2002).

It nevertheless seems likely, however, that tropical insect communities may show patterns similar to other organisms, with competition predominantly influencing community composition in wet forest while in dry seasonal forests, habitat filtering due to stronger environmental seasonality, is a major community assembly driver. This might be expected because of the hypothesis of biotic interactions being stronger in more diverse communities and weaker in less diverse communities (Schemske *et al.* 2009). Moreover, the intensity or frequency of biotic interactions is likely to vary within tropical forests with varying degrees of seasonality. For example, Coley & Barone (1996) found higher rates of leaf herbivory occuring in wet forests compared to dry forests.

Vertical strata within rainforests offer much ‘steeper’ climatic gradients compared to those found across latitude and elevation dimensions (Scheffers *et al.* 2013). Indeed, abiotic conditions shift progressively from the understory to the canopy, particularly in terms of temperature, humidity and light (Johansson 1974, Fetcher *et al.* 1985), with canopy conditions within lowland forests exhibiting warmer and drier conditions compared to the understory. The varying abiotic environment is accompanied by a conspicuous change in the biological communities inhabiting the forest strata. This is the case for Neotropical butterfly assemblages, which show clear stratification patterns in terms of abundance and composition (DeVries & Walla 2001, Fermon *et al.* 2003, Fordyce & DeVries 2016 and citation therein).

As a consequence, species exhibit morphological traits potentially associated with flight height. Understory species are hypothesized to have high wing-area/thoracic-volume ratios, associated with slower flight (Hall & Willmott 2000), and higher wing aspect ratio (*e.g.*, long and narrow wings) and wing loading (Mena & Checa in prep.). Wings with high aspect ratio are aerodynamically more efficient for gliding (Betts & Wooton 1988), a flight mode adapted to lower temperatures in the understory as it is energetically less expensive (Ellington 1985). It is less clear how these traits vary across sites with varying seasonality, but it is likely that lower aspect ratio values occur in species inhabiting the wet forest compared to those within dry forests, since wet forest butterfly species tend to be rarer, more territorial and display perching behavior to find mates (see Rutowski 1991, Hall & Willmott 2000), traits more frequent in more diverse, competitive environments.

The goal of the present study is, therefore, to test the hypothesis that assembly processes that limit species similarity predominantly occur in more ‘stable’ abiotic environments, whereas habitat filtering is a major driver of community composition within more variable environments at regional (*e.g.*, aseasonal vs seasonal forests) and local scales (*e.g.*, understory vs. canopy). In other words, I expect the importance of competition as an assembly mechanism to decrease from the wet forest to dry forest, and at local scales from the understory to the canopy, whereas the contrary pattern is expected to occur for habitat filtering, being prevalent in the dry forest and understory strata. A combined approach of phylogenetic- and trait-based analyses using forewing length and aspect ratio as traits, were used to test these hypotheses.

## Methods

### Census Techniques

Butterflies were sampled using bait traps at three study sties: Canande River Reserve (CR, wet forest), Lalo Loor Dry Forest (LLDFR, transition forest) and Jorupe Reserve (JR, dry forest). Sampling was performed every 2 months (*e.g.*, six times a year) over 7 days each sampling period for three consecutive years from November 2010 to September 2013. Van Someren-Rydon bait traps were used with two different types of baits: rotting prawn fermented for 13-18 days, and banana fermented for 2 days. Traps were checked daily during the first 7 days of the sampling month from 9 am to 3 pm; traps were opened and baited on the first trapping day, and checked over the next 6 days with trapped butterflies being collected, except for the most abundant species, which were marked and released.

Two transects were established at each reserve with each transect containing eight sampling positions. Two baited traps were set up at each sampling position in two different strata, understory (1.5 meters above the ground) and canopy (10-25 m depending on the ecosystem sampled). The use of banana and shrimp baits alternated between positions, thus neighboring positions had different types of bait. Canopy and understory traps in the same position were baited with the same type of bait. All collected material was examined and identified to species, and classified following the higher taxonomic classification of Wahlberg *et al.* (2009). Legs from dried specimens were detached and stored in vials for subsequent DNA analyses.

### DNA extraction and phylogenetic analyses

DNA extraction was done from 1-2 legs (depending on butterfly size) of dried specimens using the Qiagen DNEasy extraction kit. The ‘barcode’ section of the mitochondrial gene *cytochrome oxidase 1* (COI) was sequenced for all Nymphalidae species collected across study sites, only nymphalids were included as the DNA extraction and amplification procedures were better understood for these species, and owing to resource constraints to sequence more species. Phylogenetic analyses therefore only included this taxonomic group. Sequences of two species, *Hamadryas arinome* and *Nessaea aglaura*, were obtained from GenBank. Additionally, sequences of the following species were provided by colleagues currently researching these taxa: *Memphis artacaena, M. aureola, M. cleomestra, M. mora, M. nenia, Archaeoprepona amphimachus, Prepona gnorima, P. werneri*, *Junonia genoveva* and *Marpesia chiron*.

The phylogeny based on COI sequences was constrained by the generic level phylogeny of Wahlberg *et al.* (2009) to confidently resolve deeper nodes. In order to quantify morphological traits, one photograph per individual was taken and the software ImageJ imaging was used to carry out measurements on the digital photographs. Ten individuals per species were measured and traits averaged, except in cases where species abundances was <10, where only 1-2 specimens were measured. Only males were measured owing to low captures of females with baited-traps, as well as to females’ higher morphological variability (DeVries *et al.* 2010). Variables measured included forewing length, measured from the base to the apex, and aspect ratio (AR), calculated as 4*forewing length^2*^ wing area^−1^ (Betts & Wooton 1988, Dudley 1990). Forewing length was used as a proxy of body size, as previous research found it is a robust proxy (Shahabuddin & Ponte, 2005, Ribeiro & Freitas 2011).

### Statistical Analyses

Blomberg’s K statistic was used to measure the phylogenetic signal in morphological tratis, AR and FWL (Blomberg *et al.* 2003), based on the species-level phylogenetic tree. This statistic estimates the trait variability within a phylogeny, with K>1 meaning more trait conservatism than expected under a Brownian model of evolution, in contrast to more trait convergence than expected under the same model of evolution when K<1. Significance of Blomberg’s K statistic was assessed generating 999 random combinations of trait values by shuffling the traits across the tips of the phylogeny 999 times; the observed k statistic was further compared to the distribution of k values generated by the null model. Trait conservatism exists if observed K values fall in the upper 2.5% of the null distribution (Kraft *et al.* 2010).

### Phylogeny-based analyses

Phylogenetic analyses were performed using the constrained species-level phylogenetic tree. Phylogenetic structure of butterfly communities was estimated using two indices: the Net Relatedness Index (NRI) and the Nearest Taxon Index (NTI). These metrics allow determining whether communities include a random set of species from the regional pool or, by contrast, a deterministic process drives community composition (*e.g.*, phylogenetic clustering or even dispersion is observed). NRI refers to the standardized effect size of the mean phylogenetic distance (MPD) across species in the local communities, and NTI corresponds to the standardized effect size of the mean nearest taxon distance (MNTD) (Donoso 2013). Positive values of NTI and NRI denote phylogenetic clustering, whereas negative values indicate communities that are phylogenetically evenly dispersed. Standardized effect sizes were obtained by comparing observed values to those generated by null models, standardized by the standard deviation of the null distribution. Two null models were used, ‘Taxa Labels’ which generate random communities by shuffling the tips of the phylogeny, and ‘Sample Pools’ that creates null communities by drawing species from the sample pool. Communities were considered significantly clustered or evenly dispersed if the observed distance of MPD and MNTD was above or below 2.5% of those within the null distributions. In addition, higher NTI values with respect to NRI mean more clustering at the tips of the phylogeny, rather than tree-wise.

### Trait-based Analyses

Following the approach developed by Kraft *et al.* (2007) and Kraft & Ackerly (2010), we assessed assembly mechanisms of butterfly communities by analyzing trait dispersion within these communities. we estimated several indices sensitive to niche differentiation and habitat filtering. First, we calculated the indices ‘range’ and ‘variance’ of traits, which are expected to decrease within a community in the presence of an environmental filter. In addition, three indices sensitive to trait dispersion were estimated: standard deviation of nearest neighbor distance along trait axes divided by trait range (SDNNr), standard deviation of successive neighbor distances along trait axes divided by trait range (SDNDr), and kurtosis. The standard deviation of NN and ND is an index representing how regularly spaced are species along the trait axis, whereas division by range is used to partially correct for effects of habitat filtering, thus detecting niche differentiation in an environment of ecological filters. Moreover, both SDNNr and SDNDr are estimated by detecting the most similar species to each successive species within the assemblage. In communities where biotic interactions act as assembly mechanisms, the spacing of trait values remains constant, and as a consequence kurtosis, SDNDr and SDNNr are expected to decrease. One-tailed Wilcoxon tests were used to determine whether these metrics significantly differed from zero. All phylogenetic analyses were performed using the R package ‘Picante’ (Kembel *et al.* 2010).

## Results

A total of 6466 specimens representing 142 species of Nymphalidae were recorded from November 2010 to September 2013 within wet, transition and dry forests from western Ecuador. Observed species richness decreased gradually from wet forests (98 species), to transition (60) and dry forests (32) in the southern part. Few species dominated the samples; in particular, within dry forest butterfly communities, two species, *Fountainea ryphea* and *Hamadryas amphichloe*, comprised 65% of specimens collected (1518 and 767 individuals, respectively). Species abundances were more even within the wet forest since the most abundant species, *Memphis cleomestra* and *Hamadryas amphinome*, were represented by 158 and 143 individuals, respectively, which corresponded to 23% of the total sample.

### Phylogenetical Signal In Traits

Butterfly species significantly differed in terms of body size and wing aspect ratio across study sites (Fig. 1). The former gradually decreased from wet forest to dry forests with butterfly species’ forewing length varying in mean values from 35.73 mm to 32.48 mm, respectively; whereas this trait averaged 35.50 mm in the transition forests. In terms of AR, butterfly species from the transition forests presented the highest mean value of 5.78, followed by dry forest species (5.71) and wet forest species (5.65). Hence, butterflies inhabiting dry forests on average tended to have smaller body size (lower values of FWL) and longer, narrower wings (high AR) compared to those from the other study sites. Wilcoxon tests revealed that the standardized mean values of both traits highly significantly differed among butterfly communities across ecosystems. Moreover, Blomberg’s K values for AR and FWL revealed that traits were more convergent than expected under the Brownian model of evolution (k= 0.20 p= 0.00; k= 0.18 p= 0.00, respectively), but more conserved than predicted by a random association of traits and phylogeny as shown by their associated p values (all p values < 0.05). Hence, owing to p values less than 0.05, traits were considered as conserved (showing phylogenetic signal) for the interpretation of results, with clustered patterns of phylogenetic relatedeness indicating the mechanisms limiting similarity of species are shaping community composition. By contrast, even phylogenetic dispersion indicated ecological filters as assembly mechanisms. K values also showed that AR is more evolutionarily labile, while forewing length was more conserved.

**Figure 1.**
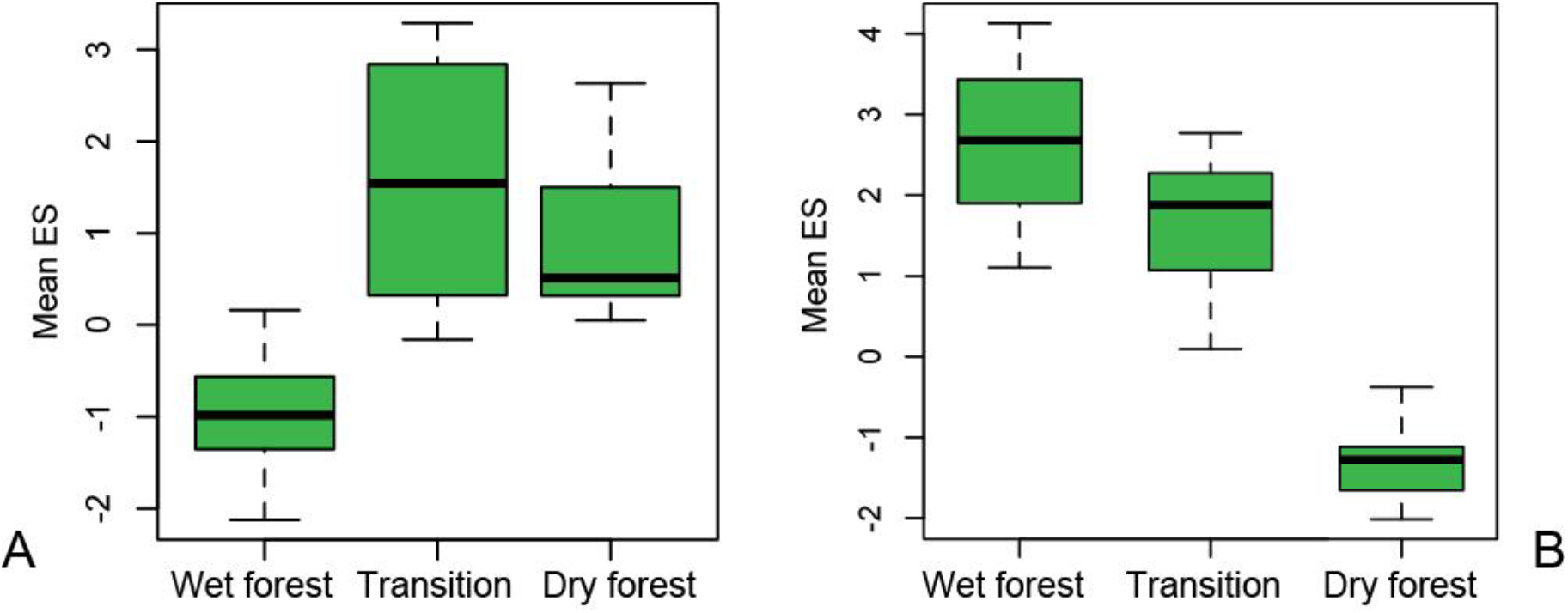
Box plots of standardized mean values of species traits within communities from wet, transition and dry forests; green color means highly significant differences were found among sites (p< 0.01) according to Wilcoxon Tests. A) Box plot corresponding to wing Aspect Ratio. B) Box plot corresponding to forewing length.

### Phylogenetic-based Tests Of Community Composition

Phylogenetic-based tests revealed significant levels of phylogenetic clustering (Fig. 2). According to NRI values, the wet forest community was composed of more phylogenetically closely related species than expected, regardless of the null model used, namely by shuffling the tips of the phylogeny (Taxa Label, TL) or drawing species from the sample pool (Sample Pool, SP). (Fig. 2). Likewise, NTI values revealed phylogenetic clustering in dry forest communities, but solely for SP null models when considering NRI (NRI= 0.3 p= 0.00). The distribution of observed NRI and NTI values for wet and dry forest communities was significantly shifted above null expectation. Species within butterfly communities from transition forest were random with respect to phylogeny (non-significance detected in both NRI and NTI indices).

**Figure 2.**
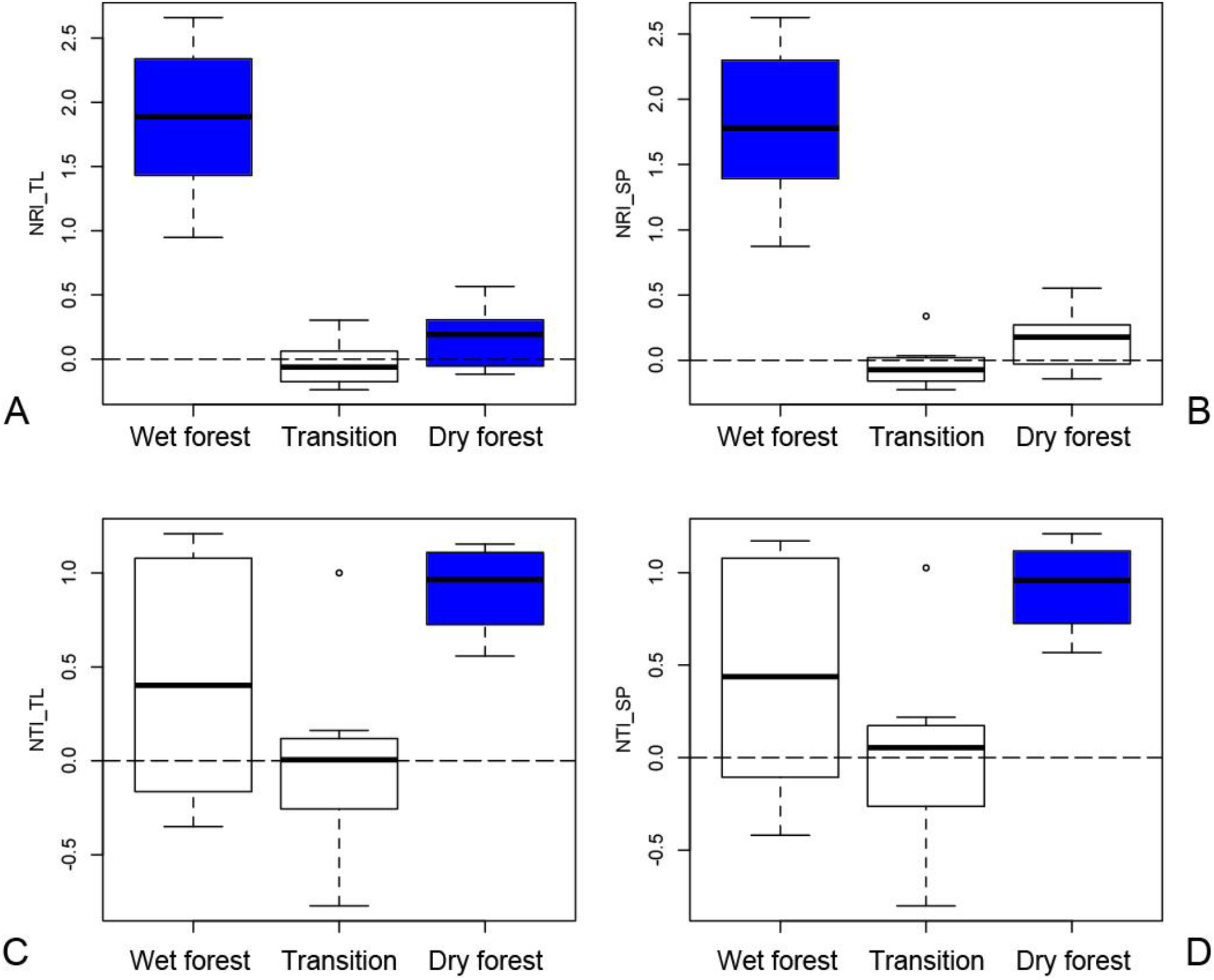
Indices of phylogenetic community structure, Net Relatedness Index (NRI) and Nearest Taxon Index (NTI) estimated using different null models for butterfly communities within wet, transition and dry forests; blue color means taxa are more related than expected or phylogenetically clustered. A) NRI using Taxa Label null model. B) NRI using Sample pool null model. C) NTI using Taxa Label null model. B) NTI using Sample pool null model.

In terms of phylogenetic structure of communities across strata within sites, similar patterns of phylogenetic clustering were detected. In cases using Taxa Label null models, co-occurring taxa from the canopy and understory communities within wet forest tended to be more closely related than expected based on values of NRI and NTI (Table 1). Moreover, phylogenetic clustering was detected for canopy (NTI= 0.98 p<0.01) and understory communities (NTI= 1.38 p<0.01) distributed in the dry forest. With respect to transition forests, species within canopy communities were random or phylogenetically more related than expected (NTI= 0.43 p>0.05, NTI= 0.26 p< 0.05), whereas species in understory communities were random (NRI= −0.47 p> 0.05, NTI= −0.36 p>0.05) (Table 1). Similar trends were revealed when using the Sample Pool algorithm, namely by drawing species from the sample pool to generate null communities.

**Table 1.** Indices of phylogenetic community structure, Net Relatedness Index (NRI) and Nearest Taxon Index (NTI) estimated using different null models (*i.e.*, Taxa Label TL or Sample Pool SP) for butterfly communities within wet, transition and dry forests; asterisk or doubled asterisks mean observed indices were significantly or highly significantly different from expected values estimated with null models.

|  | NRI_TL | NRI_SP | NTI_TL | NTI_SP |
| --- | --- | --- | --- | --- |
| Wet forest |  |  |  |  |
| Canopy | 1.64** | 1.70** | 0.46* | 0.44* |
| Understory | 2.32** | 2.15** | 0.87* | 0.77* |
| Transition |  |  |  |  |
| Canopy | 0.26* | 0.29* | 0.43 | 0.43 |
| Understory | -0.47 | -0.50 | -0.36 | -0.37 |
| Dry forest |  |  |  |  |
| Canopy | 0.29 | 0.30 | 0.98** | 0.97** |
| Understory | -0.13 | -0.10 | 1.38** | 1.39** |

### Trait-based Tests of Community Composition

Non-random patterns of even spacing with respect to AR were solely detected for butterfly community composition within transition forests, with two metrics sensitive to niche differentiation showing a significant reduction for AR: SDNNr (p= 0.01) and SDNDr (p= 0.02) (Fig. 3). All other metrics including variance, range and kurtosis were not significantly reduced relative to the null expectation for any study site, revealing random patterns of AR distribution.

**Figure 3:**
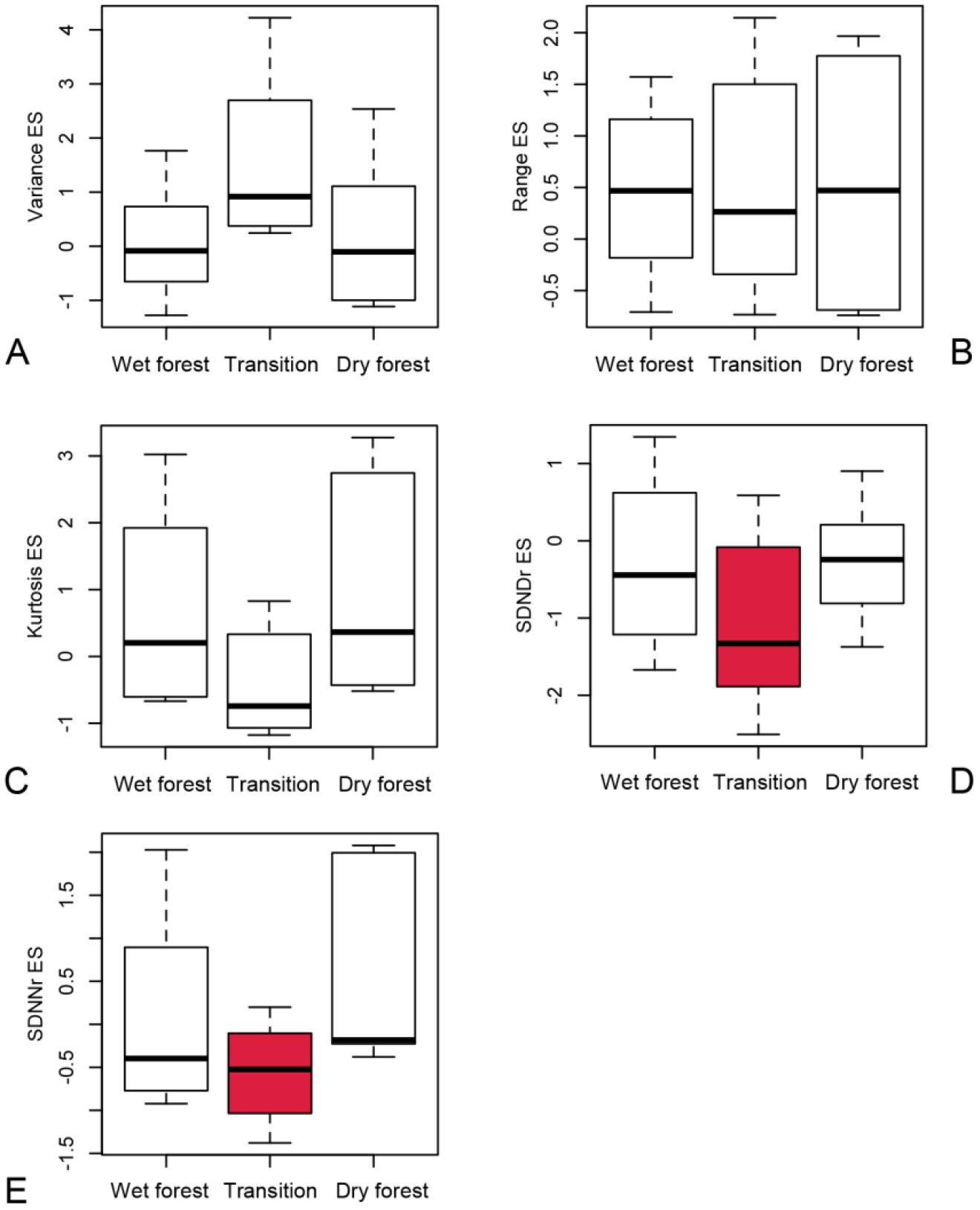
Standardized values of metrics estimated for wing aspect ratio of butterfly communities from wet, transition and dry forests; metrics are sensitive to habitat filtering (A-B) and niche differentiation (C-E). Red color means indices significantly decreased with respect to their null distribution according to Wilcoxon tests (p<0.05) showing evidence for competition. A) Variance. B) Range. C) Kurtosis. D) Standard deviation of successive neighbor distances along trait axes divided by range (SDNDr). C) Standard deviation of nearest-neighbor distance along trait axes divided by range (SDNNr).

Trait-based tests using body size detected nonrandom patterns solely within dry forests as SDNDr and kurtosis were highly significantly lower relative to the null expectation, revealing even dispersion within this community (Fig. 4). There was no significant decrease in range and variance in traits within wet forest communities (p= 0.054 and p= 0.074, respectively), although p values close to 0.05 suggested a trend of phylogenetic clustering. Random patterns of body size distribution were detected for transition forest communities, wet forest communities in relation to metrics sensitive to niche differentiation, and dry forest communities with respect to variance and range.

**Figure 4:**
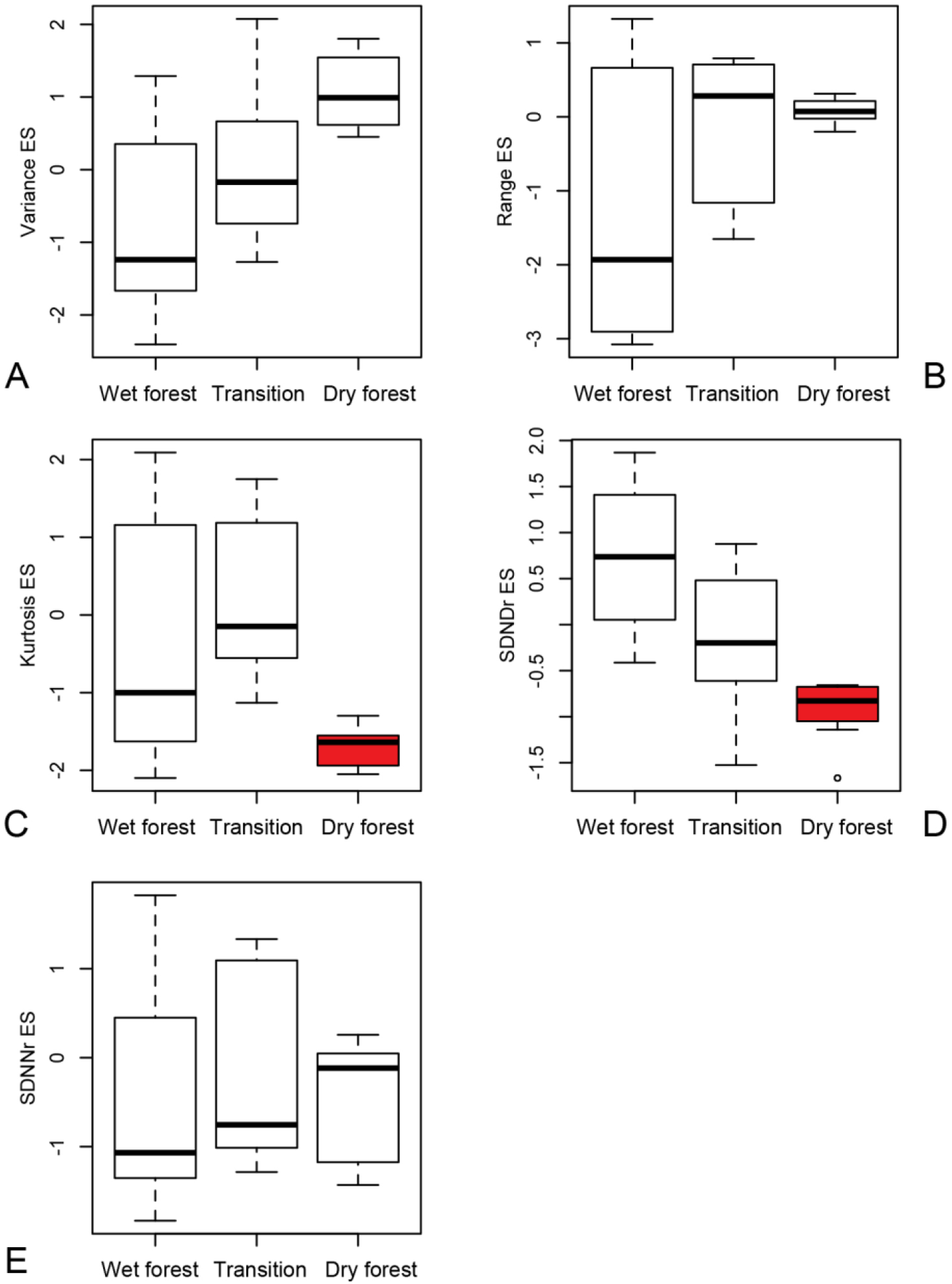
Standardized values of metrics estimated for forewing length of butterfly communities from wet, transition and dry forests; metrics are sensitive to habitat filtering (A-B) and niche differentiation (C-E). Blue and red color mean indices significantly decreased with respect to their null distribution according to Wilcoxon tests (p<0.05) showing evidence for habitat filtering and competition, respectively. A) Variance. B) Range. C) Kurtosis. D) Standard deviation of successive neighbor distances along trait axes divided by range (SDNDr). C) Standard deviation of nearest-neighbor distance along trait axes divided by range (SDNNr).

Along the vertical forest dimension, similar patterns of trait even spacing were detected for dry forest communities within the canopy and understory. With respect to AR, even dispersion was detected since SDNDr significantly decreased within understory and canopy communities, along with kurtosis for understory assemblages (Table 2).

**Table 2.** Standardized values of metrics estimated for wing aspect ratio of butterfly communities across strata within wet, transition and dry forests; metrics are sensitive to habitat filtering (variance and range) and niche differentiation (kurtosis, SDNDr and SDNNr). SDNDr stands for standard deviation of successive neighbor distances along trait axes divided by range, whereas SDNNr refers to standard deviation of nearest-neighbor distance along trait axes divided by range. Asterisk or asterisks refer to indices that significantly decreased with respect to their null distribution according to Wilcoxon tests (p<0.05 and p<0.01, respectively).

|  |  | Range | Variance | Kurtosis | SDNNr | SDNDr |
| --- | --- | --- | --- | --- | --- | --- |
| Wet forest | Canopy | 0.9 | 0.25 | 0.6 | 0.5 | -0.4 |
|  | Understory | -0.3 | -0.9* | 0.2 | 0.35 | -0.25 |
| Transition | Canopy | 0.1 | 1.1 | -0.3 | -0.15 | -0.55 |
|  | Understory | 0.9 | 0.75 | -0.75 | -0.9 | -0.3 |
| Dry forest | Canopy | -0.5 | 0.75 | -0.25 | -0.6 | -0.5* |
|  | Understory | 0.8 | 1 | -0.9** | 0.85 | -0.56* |

Likewise, all metrics sensitive to niche partitioning significantly decreased within understory and canopy communities for body size analyses (Table 3). Analyses focused on body size additionally revealed even dispersion within the understory from wet forest communities (*e.g.*, kurtosis and SDNNr significantly decreased), along with a clustering pattern within the canopy community (*i.e.*, range and variance significantly decreased). On the other hand, metrics based on AR also revealed a clustering pattern in the understory community within wet forest communities (variance decreased compared to null expectation, Table 3). Summaries of the outcomes from phylogeny- and trait-based analyses are shown in Table 4, 5 and 6.

**Table 3.**
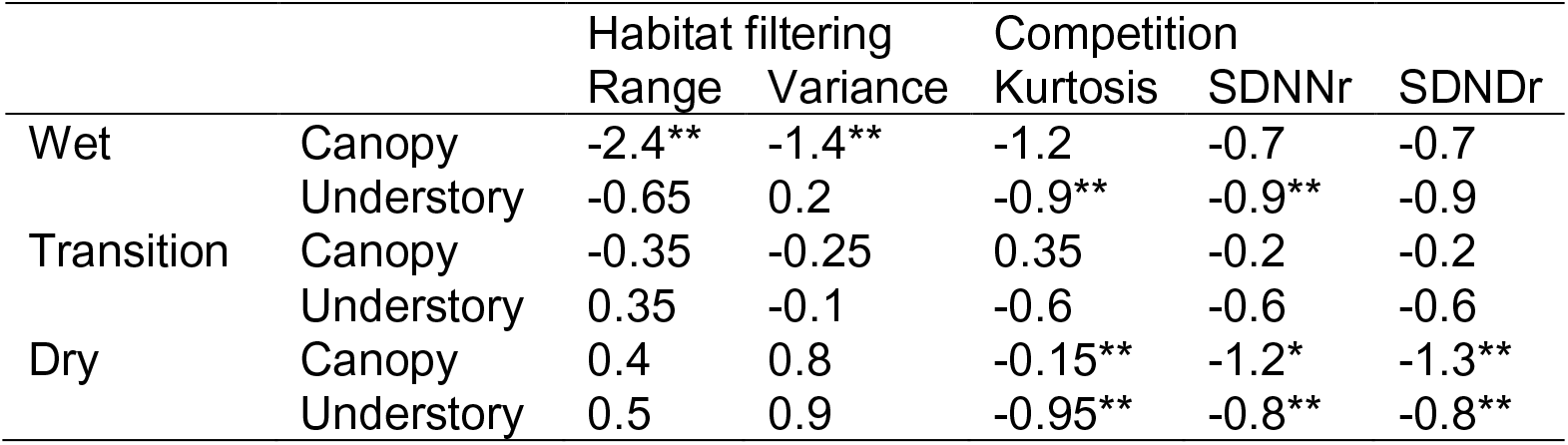
Standardized values of metrics estimated for forewing length of butterfly communities across strata within wet, transition and dry forests; metrics are sensitive to habitat filtering (variance and range) and niche differentiation (kurtosis, SDNDr and SDNNr). Asterisk or asterisks refer to indices that significantly decreased with respect to their null distribution according to Wilcoxon tests (p<0.05 and p<0.01, respectively).

|  |  | Habitat filtering |  | Competition |  |  |
| --- | --- | --- | --- | --- | --- | --- |
|  |  | Range | Variance | Kurtosis | SDNNr | SDNdr |
| Wet | Canopy | -2.4** | -1.4** | -1.2 | -0.7 | -0.7 |
|  | Understory | -0.65 | 0.2 | -0.9** | -0.9** | -0.9 |
| Transition | Canopy | -0.35 | -0.25 | 0.35 | -0.2 | -0.2 |
|  | Understory | 0.35 | -0.1 | -0.6 | -0.6 | -0.6 |
| Dry | Canopy | 0.4 | 0.8 | -0.15** | -1.2* | -1.3** |
|  | Understory | 0.5 | 0.9 | -0.95** | -0.8** | -0.8** |

**Table 4.**
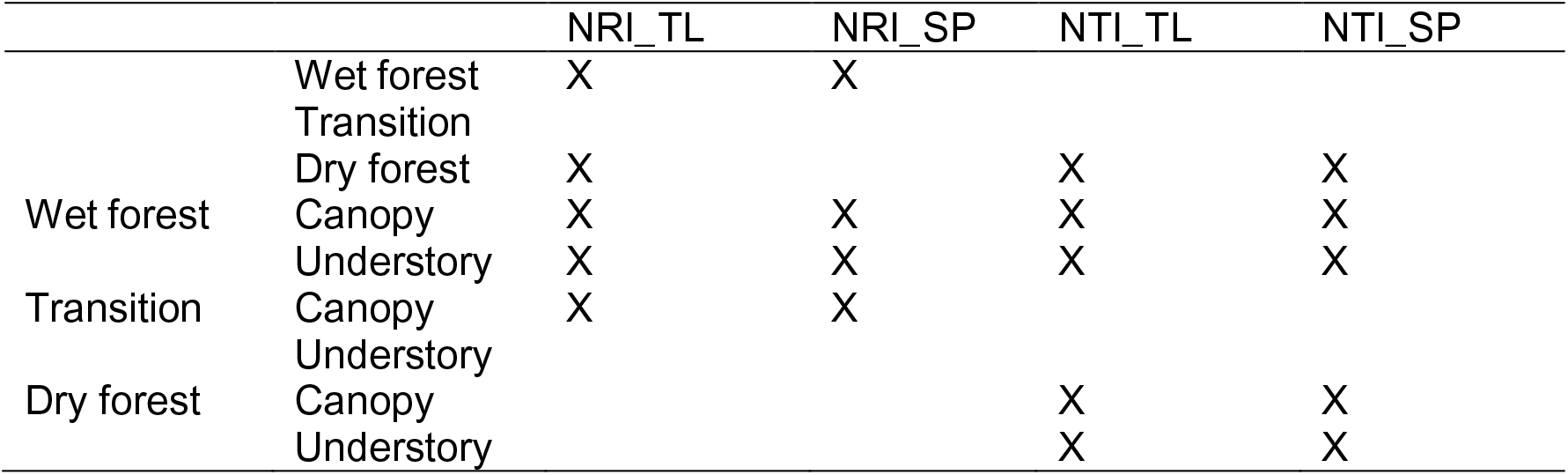
Summary of outcomes from phylogenetic-based analyses based on indices of phylogenetic community structure, Net Relatedness Index (NRI) and Nearest Taxon Index (NTI) estimated using different null models (*i.e.*, Taxa Label TL or Sample Pool SP) for butterfly communities within wet, transition and dry forests. The letter X denotes habitat filtering was detected as a prevalent assembly mechanism within these butterfly communities. Blank cells indicate random patterns of species distribution were found.

**Table 5.** Summary of outcomes from trait-based analyses based on the standardized values of metrics estimated for body size and aspect ratio (AR) of butterfly communities within wet, transition and dry forests; metrics are sensitive to habitat filtering (variance and range) and niche differentiation (kurtosis, SDNDr and SDNNr). The letter X denotes which assembly mechanism was detected as significant for each butterfly community. Blank cells indicate random patterns of trait distribution within communities were found.

| Body size | Habitat Range | filtering Variance | Competition Kurtosis | SDNNr | SDNDr |
| --- | --- | --- | --- | --- | --- |
| Wet forest |  |  |  |  |  |
| Transition |  |  |  | X | X |
| Dry forest |  |  |  |  |  |
| AR |  |  |  |  |  |
| Wet forest |  |  |  |  |  |
| Transition |  |  |  |  |  |
| Dry forest |  |  | X |  | X |

**Table 6.**
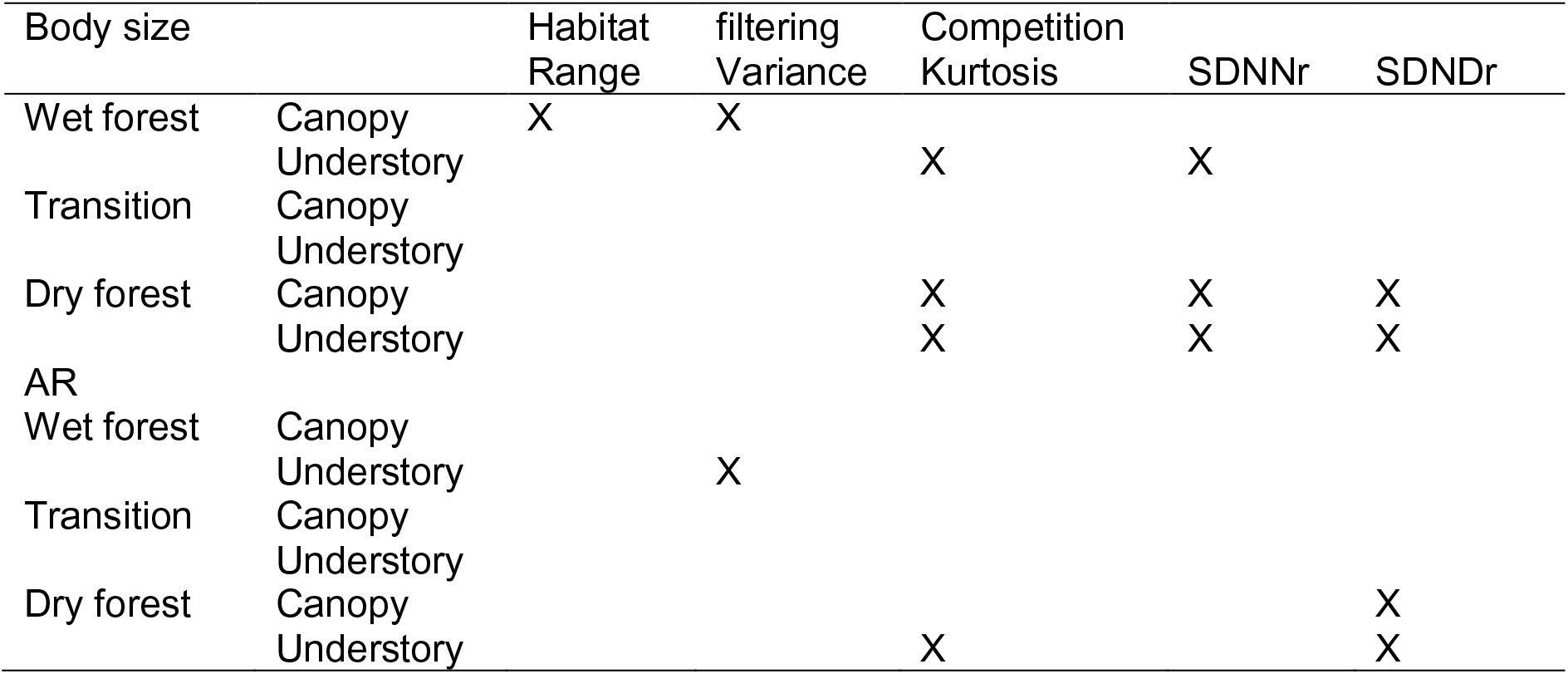
Summary of outcomes from trait-based analyses based on the standardized values of metrics estimated for body size and aspect ratio (AR) of butterfly communities across strata within wet, transition and dry forests; metrics are sensitive to habitat filtering (variance and range) and niche differentiation (kurtosis, SDNDr and SDNNr). The letter X denotes which assembly mechanism was detected as significant for each butterfly community. Blank cells indicate random patterns of trait distribution within communities were found.

## Discussion

Based on the results of the present study, I rejected the hypothesis that assembly processes that limit species similarity (*i.e*., competition) are likely to predominantly occur in more ‘stable’ abiotic environments, whereas habitat filtering can be a major driver of community composition within more variable environments at regional (*i.e.*, aseasonal vs seasonal forests) and local scales (*i.e.*, understory vs. canopy). My study of assembly mechanisms revealed the opposite pattern, with stronger evidence for the action of ecological filters in the assembly of butterfly communities from the wet aseasonal forests, and competition likely to be a major assembly process within dry seasonal forests.

Phylogenetic- and trait-based analyses revealed mostly non-random patterns of phylogenetic structure within butterfly communities along local and regional environmental gradients in western Ecuador. I found both traits AR and body size were evenly dispersed within communities from the dry forest even at local scales across strata communities, which is consistent with the competition hypothesis. Additionally, phylogenetic tests revealed co-occurring species were more closely related than expected by chance within canopy and understory communities from the dry forest, pattern consistent with the habitat filtering hypothesis. Similarly, habitat filtering was found as a significant assembly mechanism within both the canopy and understory communities from wet forests, with less evidence of trait even dispersion (*i.e.*, competition) along this gradient (*i.e.*, significant outcomes solely from body size analyses). With respect to transition forest communities, random patterns of phylogenetic structure were mostly observed, except for local scale analyses that revealed phylogenetic clustering within canopy butterfly assemblages and trait-based analyses revealing even dispersion of AR, results consistent with habitat filtering and competition, respectively.

These general conclusions do not mean that competition or habitat filtering are mutually exclusive depending on the ecosystem analyzed, or that other factors (*e.g.*, evolutionary) have affected community composition, only that at the scale of the analyses presented here and the approach used (Kraft *et al.* 2007), the role of one mechanism might be prevalent above the effect of the other (see Graham *et al.* 2009). Indeed, other biotic interactions such as mutualistic mimicry have been found to shape Andean butterfly communities (Chazot *et al.* 2014). Furthermore, predation (see Schulze *et al.* 2001) and distribution of hostplants (see Beccaloni 1997, Willmott & Mallet 2004) can also structure butterfly community composition across strata within tropical forests.

Our understanding of the importance of competition in tropical butterfly communities is very limited (Grøtan *et al.* 2012). The results found here confirm that it is challenging to predict how competition can vary along environmental gradients where other processes to prevent it commonly occur, such as niche partitioning in time *(i.e.,* seasonality). It is likely that, indeed, that the latter prevents competition being a prevalent assembly mechanism within wet forests, in contrast to what was expected of this mechanism shaping butterfly communities in more ‘benign’ aseasonal environments.

Temporal species turnover has been reported to occur in tropical butterfly communities from aseasonal forests (Checa *et al.* 2009) with community similarity displaying annual cycles that peak in the driest months (Valtonen *et al.* 2013, Grotan *et al.* 2014). It is likely that the incidence of drought throughout 4-6 months strongly constrains the ‘time’ resource within dry forests, which can further impose a challenge for species survival thereby increasing competition within communities, explaining the prevalence of competition as an assembly mechanism. However, other factors might come into play in order to explain my results (see below).

Most stabilizing assembly mechanisms, including competition and habitat filtering, are not mutually exclusive; in fact, these processes can generate interacting effects on biological communities (Kraft *et al.* 2007, Kraft *et al.* 2010). As a consequence, trait or phylogenetic distribution is altered and random patterns can occur, as suggested by Swenson & Enquist (2009) for a research study focused on tropical plant communities within dry forests, where the simultaneous occurrence of clustering and even dispersion of traits led to random phylogenetic community structure. As a consequence, it is not possible to disentangle whether the random patterns of trait and phylogenetic distribution found here, particularly for wet and transition forests (see summary Table 4-5, 4-6), are indeed showing equalizing mechanisms (the most remarkable example being Hubell’s neutral theory) as assembly processes within these communities, or by contrast, are the result of interacting effects of competition and habitat filtering.

Furthermore, outcomes derived from trait-based analyses might depend on studied traits (Kraft *et al.* 2010). For example, Sedio *et al.* (2012) found that hydraulic but not photosynthetic traits largely explained phylogenetic clustering within species communities of the hyperdiverse plant genus *Psychotria* in Panama. Hence, further research incorporating other functional traits of butterflies into analyses of community structure could provide more evidence to patterns described in the present study. In addition, including more taxonomic groups (not only Nymphalidae) into the analyses could further contribute towards our understanding as it will increase the phylogenetic diversity present in the local and regional communities, as well as an increased range of trait variation.

An interesting result is that butterfly species are using the traits of AR and body size as adaptations to different processes depending on the ecosystem and strata analyzed. For example, niche partitioning involved both traits within understory and canopy communities from dry forests, whereas solely body size remained relevant for competition within the wet forest (understory) and AR within the transition forest. On the other hand, body size and AR are relevant adaptations for the abiotic environment acting as ecological filters in wet forests within the canopy and understorey, respectively.

Furthermore, a trend existed of smaller body size and higher AR (*i.e.*, long and narrow wings) in butterfly species inhabiting dry forest compared to wet forest. The genus *Heliconius* showed the highest AR values in the present study, and population studies about this well-known genus in the tropics revealed competition occurs for larval resources for some species but not others (Rodrigues & Moreira 2004) as populations are below carrying capacity of hostplants (Ehrlich & Gilbert 1973) and niche partitioning occurs among instars (*e.g.,* younger instars feed on apical and new shoots) (Rodrigues & Pires 1999). Intra-specific competition for mates can also affect population dynamics of *Heliconius* (see Delnert *et al.* 1994).

These results indicated that morphological traits, namely body size and AR, are likely involved in competition among Neotropical butterfly species. Moreover, the results presented here provide insights into assembly mechanisms in one of the richest butterfly faunas worldwide, revealing competition along with ecological filters as significant drivers of community composition.

